# Genetics of Cocaine Consumption and Preference in *Drosophila melanogaster*

**DOI:** 10.64898/2026.08.20.746013

**Authors:** Jeffrey S. Hatfield, Vijay Shankar, Robert R. H. Anholt, Trudy F. C. Mackay

**Author notes:** Co-Corresponding authors: Contact Information: Dr. Trudy F. C Mackay, Institute for Human Genetics, 114 Gregor Mendel Circle, Greenwood, SC 29646, Dr. Robert R. H. Anholt, Institute for Human Genetics, 114 Gregor Mendel Circle, Greenwood, SC 29646.

## Abstract

Cocaine Use Disorder (CUD) poses a significant public health and socioeconomic challenge. Determining the genetic basis of predisposition for development of CUD is challenging in human populations but can be studied in *Drosophila*. We assessed cocaine consumption and cocaine preference of 74,875 flies from 598 sequenced, wild-derived, inbred lines from the expanded *Drosophila melanogaster* Genetic Reference Panel (DGRP3). We found significant genetic variation, sexual dimorphism, and genetic variation in sexual dimorphism for these traits. Whereas most lines showed cocaine avoidance, ∼10% of the lines showed innate cocaine preference in at least one sex. Genome-wide association analyses for cocaine consumption, preference, and micro-environmental variance of these traits identified 2,155 polymorphisms in/near 866 genes that were enriched for Gene Ontology terms associated with neurogenesis, development, and behavior. Many of the associated genes had human orthologs with known associations with CUD and other substance use disorders as well as psychiatric and behavioral traits. We confirmed causal associations with cocaine preference for three polymorphisms with large effect sizes by assessing their effects in DGRP3 lines not included in the initial association analyses. Pairwise associations between these polymorphisms exhibited suppressing epistasis. These polymorphisms are in genes with human orthologs that fulfill essential functions in the nervous system, including the glucose transporter *SLC2A8*; *KCNC2*, a subunit of the voltage gated potassium channel; and *CHRNA7*, a nicotinic cholinergic receptor subunit. Thus, studies on *Drosophila* can provide insights into the genetic and neural mechanisms of CUD.

## Introduction

Cocaine Use Disorder (CUD) is a significant public health and socioeconomic problem. Cocaine contributes to ∼40% of drug-related emergency room visits, and incidence rates of cocaine overdose in the United States have quadrupled from 2015 (6,784) to 2023 (29,449) (NIDA https://nida.nih.gov/research-topics/trends-statistics/overdose-death-rates#Fig8). The consequences of escalating cocaine use include tachycardia, acute coronary syndrome, hyperthermia, seizures, strokes, and fatalities [1]. CUD has a substantial genetic component, with heritability estimates from human twin studies ranging from 0.39-0.44 for cocaine use, 0.32-0.79 for cocaine abuse, and 0.65-0.79 for cocaine dependence [2]. However, the success of studies mapping the genetic basis of CUD in humans has been limited by small numbers of affected individuals due to criminalization of cocaine, the inability to control environmental exposures, and comorbidities such as multi-drug use and other psychiatric disorders [2].

Only a handful of genes have been associated with cocaine dependence and related traits. Initial studies were conducted on candidate genes, which typically had small sample sizes and could not be replicated in independent samples. Exceptions are *CNR1* and the *CHRNA5-CHRNA3-CHRNB4* gene cluster [2] and a copy number variant in *NSF* [3]; variants in *CHRNA5* are also associated with crack cocaine addiction [4]. Genome wide association (GWA) analyses have found only a few associations with cocaine traits at a genome wide significance level (*P* = 5 × 10^−8^). In the first GWA study of cocaine-related traits, a genome-wide significant variant in *FAM35A* was associated with cocaine dependence; and variants in *CDK1* and *NCOR2* were associated with cocaine-induced anxiety and the number of cocaine dependence symptoms, respectively, although these variants only reached genome-wide significance in one of the populations sampled [5]. A re-analysis of these data using a gene-based test further showed an association of *NDUFB9* with cocaine dependence. A variant in *FAM78B* and an intergenic variant on chromosome 21 were genome-wide significant for cocaine dependence in a meta-analysis of the discovery and replication cohorts [6]. Another study [7] identified a genome-wide significant variant in *SLC25A16* associated with cocaine dependence, and a meta-analysis [8] identified significant association of *DRD2* and *OPRM1* with cocaine dependence. More associations were identified when more homogenous subgroups were assessed. For example, variant associations in 13 genes (*TRAK2, LINC00378, TMEM51, LPHN2, LINC01411, RP13– 20L14.1, SLC7A13, TRDN, SOGA2, RN7SL609P, SYNGR1*, *FN1, TENM3*) were significant when the population was stratified into five groups based on childhood environmental factors [9]. Four of these genes (*LINC01411*, *TRAK2*, *TMEM51, LPHN2*) were replicated in an independent sample [9]. A review of the genetic basis of the addiction, depression and anxiety (ADA) psychological disorder that focused on cocaine as the abused substance [10] identified 14 genes associated with cocaine addiction (*ABCB1, ARC, BDNF, DAT1, DRD2, GABBR1, GABRG2, GAD1, GAD2, OPRM1, PDYN, PERIOD2, SLC6A4, ZIF268*).

Each study identified different variants and genes (except for *DRD2* and *OPRM1* [8, 10]), suggesting a highly polygenic basis for cocaine-related traits. Different variants/genes were also found for different cocaine-related behaviors; and, for studies including both European American and African American ancestries, the identified variants/genes were different across ancestries. Further, the mechanism(s) by which the implicated variants affect cocaine-related behaviors remains unresolved for most of these candidate genes. Rodent studies can identify the genetic basis of variation in cocaine-related behaviors while controlling the amount and timing of cocaine exposure and reducing the co-morbidities present in human populations [11–14]. However, rodent studies are expensive and laborious.

*Drosophila melanogaster* is emerging as a powerful genetic model system with the same advantages as rodent models but with increased throughput and lower costs. Approximately 75% of human disease genes have a *Drosophila* ortholog [15]. *D. melanogaster* also has an extensive genetic toolkit enabling the identification and molecular characterization of genes and variants affecting any phenotype, including publicly available mutations and RNA interference (RNAi) constructs for most genes in the genome. The inbred lines of the *D. melanogaster* Reference Panel (DGRP) with full genome sequences are a powerful tool for GWA analyses of effects of naturally occurring variants [16–18]. The rapid decay of linkage disequilibrium with physical distance in the DGRP enables more precise mapping of candidate variants than is possible in humans or rodents.

Cocaine binds to the *Drosophila* dopamine transporter [19] and elicits impaired motor responses, excessive grooming, seizures, and mortality [20–22]. One study using the DGRP identified 726 genes associated with cocaine consumption and preference using a choice test [23]; however, this study was underpowered to detect associations that were genome-wide significant. A second study used an advanced intercross population derived from a subset of DGRP lines to identify over 214 genes with genome-wide significant variants associated with cocaine consumption in a no-choice assay [22]. Recently, the DGRP has been expanded from 205 to 1,037 inbred, sequenced lines [24]. The DGRP, version 3 (DGRP3) is a publicly available reference panel with minimal within-line heterozygosity and extensive genetic variation across the lines [24]. The DGRP3 enables evaluation of low frequency polymorphisms associated with quantitative traits, which tend to have larger effects than common polymorphisms [18]; and, if all lines are not used in the initial GWAS, permits out-of-sample validation of polymorphisms nominated in the initial GWA analysis.

We capitalized on this expanded resource and a high-throughput assay for quantifying cocaine consumption and preference in *Drosophila* [25] to perform GWA analyses for mean cocaine consumption and preference. In addition, we assessed genetic associations with within-line phenotypic variance (micro-environmental variance) [26] for cocaine-related phenotypes, identifying polymorphisms in evolutionarily conserved genes that could be relevant to variable penetrance of CUD in humans. We quantified cocaine preference and consumption for 74,875 flies in 598 DGRP3 lines. We observed significant naturally occurring genetic variation for cocaine preference and consumption, substantial sexual dimorphism for these traits, and genetic variation in sexual dimorphism. The GWA analyses implicated 2,155 polymorphisms in 866 genes associated with all cocaine-related behaviors at −*log*_10_(*P*) > 5. We functionally validated three polymorphisms associated with cocaine preference and consumption by evaluating cocaine preference in DGRP3 lines that were not included in the original GWA analysis. The evolutionarily conserved genes identified in *Drosophila* are promising targets for mechanistic/therapeutic investigation of CUD in humans.

## 2. Results

### Quantitative Genetic Analyses of Cocaine Consumption and Preference

We presented individual naive DGRP3 flies from 598 DGRP3 lines (74,875 flies total) with a choice between a sucrose control solution and a sucrose solution supplemented with 0.02% cocaine in microplate feeders [25]. After 22 hours, we quantified the cocaine consumption, sucrose consumption and cocaine preference for each fly, using absorbance values of blue dye incorporated into the solutions, adjusted for evaporation. The cocaine preference index was defined as the difference between the consumption of cocaine solution and control solution, divided by the total consumption. Therefore, positive preference values for a line indicate average consumption of more cocaine solution than control solution, while negative values indicate the reverse. We found that females on average consume more sucrose (S1 Fig., S1 Table) and cocaine than males (Figure 1A, S1 Table). However, while most lines avoid cocaine, ∼10% show preference for cocaine in at least one sex ((Figure 1B, S1 Table). On average, males demonstrate a higher preference for cocaine (mean preference index = −0.24) than females (mean preference index = −0.33) (S1 Table).

**Figure 1.**
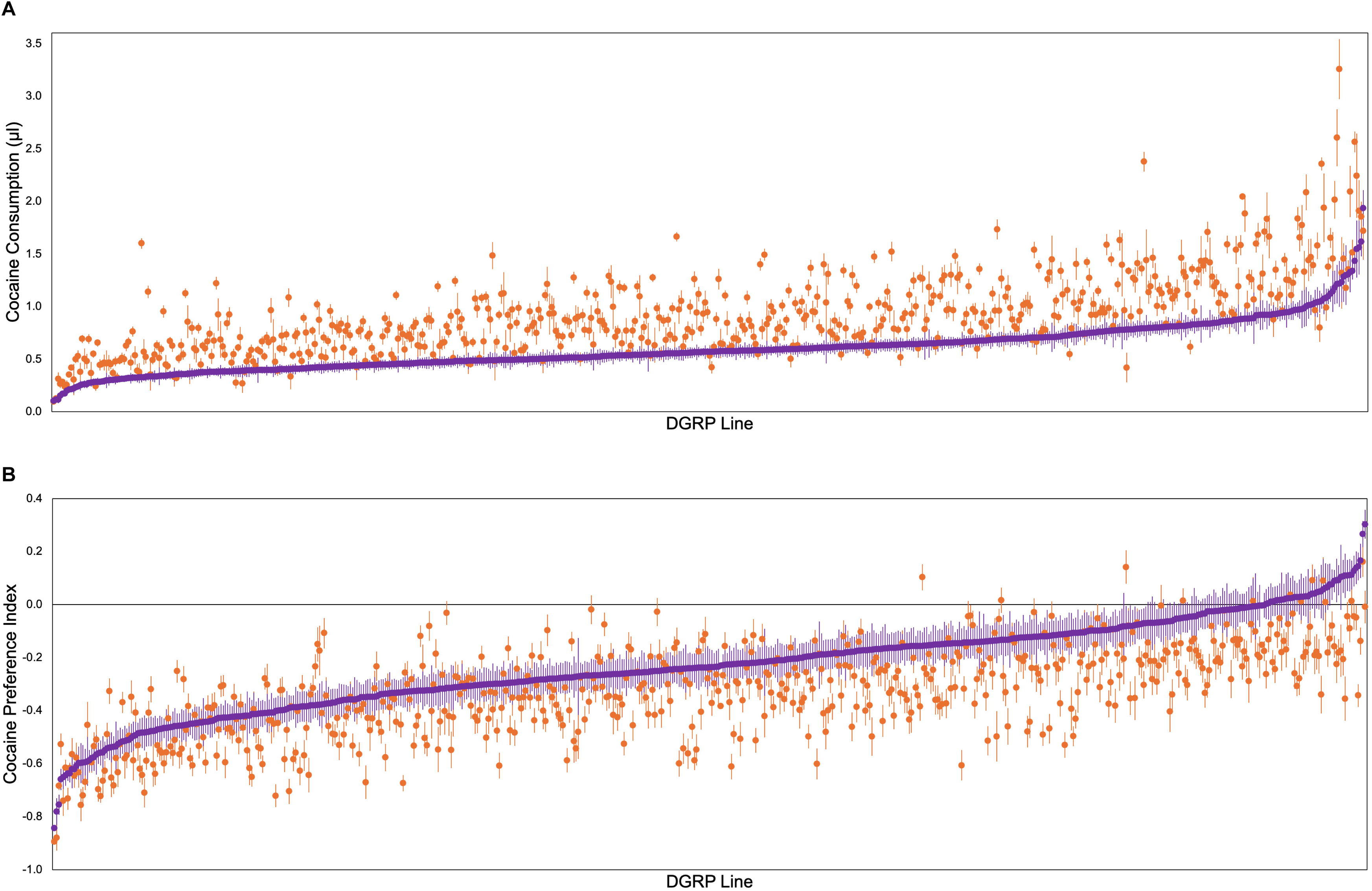
Variation in cocaine consumption and preference in the DGRP3. (A) Cocaine consumption for 598 DGRP3 lines. The circles represent line means for males (purple, *N* = 36,731) and females (orange, *N* = 38,144) and error bars are standard errors of the mean. The line means are ordered by the male cocaine consumption. (B) Cocaine preference. The mean cocaine preference index [(Mean Cocaine Consumption – Mean Sucrose Consumption)/(Mean Total Consumption)] is shown for the same 598 DGRP3 lines. The line means are ordered by the male cocaine preference index.

We performed mixed-model analyses of variance (ANOVAs) to partition variance in consumption and preference into sources of variation attributable to sex, solution (treatment), DGRP3 line, and interactions between these factors. We also ran reduced models by sex, by solution and by sex and solution (S2A Table). Overall, there is a significant effect of sex (females on average consume more than males), treatment (flies consume on average more sucrose than cocaine) and the sex × treatment interaction (the difference between sucrose and cocaine consumption is greater for females than males) (S1 Table, S2A Table). All terms including DGRP3 line are highly significant (*P* < 0.0001), indicating substantial genetic variation for consumption and preference traits. In addition, the sex × line, treatment × line and sex × treatment × line effects are significant, indicating context-dependent genetic effects for consumption traits. Broad-sense heritabilities (*H*^2^) are *H*^2^ = 0.202, *H*^2^ = 0.139, *H*^2^ = 0.239, and *H*^2^ = 0.096 for consumption pooled across sexes and treatments, cocaine consumption pooled across sexes, sucrose consumption pooled across sexes, and preference pooled across sexes, respectively (Table S2A). These individual level heritabilities are low, as expected for behavioral traits.

Given the significant genotype by sex, treatment and sex by treatment interactions, we expect cross-sex (*r_GS_*), -treatment (*r_GT_*) and -sex and treatment (*r_GST_*) genetic correlations to be less than unity [27]. Indeed, *r_GS_* = 0.735, *r_GT_* = 0.308, and *r_GST_* = 0.213 from the ANOVA of consumption pooled across sexes and treatments (S2A Table). Therefore, the genetic basis of variation in consumption of cocaine and sucrose is not identical between males and females and quite different between sucrose and cocaine. The latter observation is important because it shows that cocaine consumption and preference are not driven by common genetic variation in feeding behavior.

The DGRP3 lines are inbred; therefore, within-line variances of the consumption and preference traits primarily arise from micro-environmental plasticity [26]. We computed the environmental variance of consumption and cocaine preference for each DGRP3 line (S1 Table) and performed Levene tests for heterogeneity of environmental variance among the lines for cocaine consumption, sucrose consumption and cocaine preference, separately for males and females. All analyses are highly significant (S2B Table), indicating that there is genetic variance for micro-environmental plasticity for these traits (Figure 2, S2 Fig.). To compute the broad sense heritabilities of micro-environmental plasticity, we transformed the environmental variance estimates from each replicate microplate into *lnσ_ɛ_* values, where *σ_ɛ_* is the standard deviation of the within-line variance. This transformation is necessary because variances are not normally distributed, which violates the assumptions of ANOVA, and the *lnσ_ɛ_* metric is approximately normally distributed [26]. The broad sense heritability estimates of micro-environmental plasticity for cocaine consumption, sucrose consumption and cocaine preference are, respectively, *H*^2^ = 0.533, *H*^2^ = 0.530, and *H*^2^ = 0.404 (S2C Table). There is significant sex dimorphism for micro-environmental plasticity: females have greater micro-environmental plasticity than males for sucrose and cocaine consumption, but males have greater micro-environmental variance for cocaine preference (S1 Table, S2C Table). Genetic variation in sex dimorphism for micro-environmental plasticity is not significant for any trait (S2C Table).

**Figure 2.**
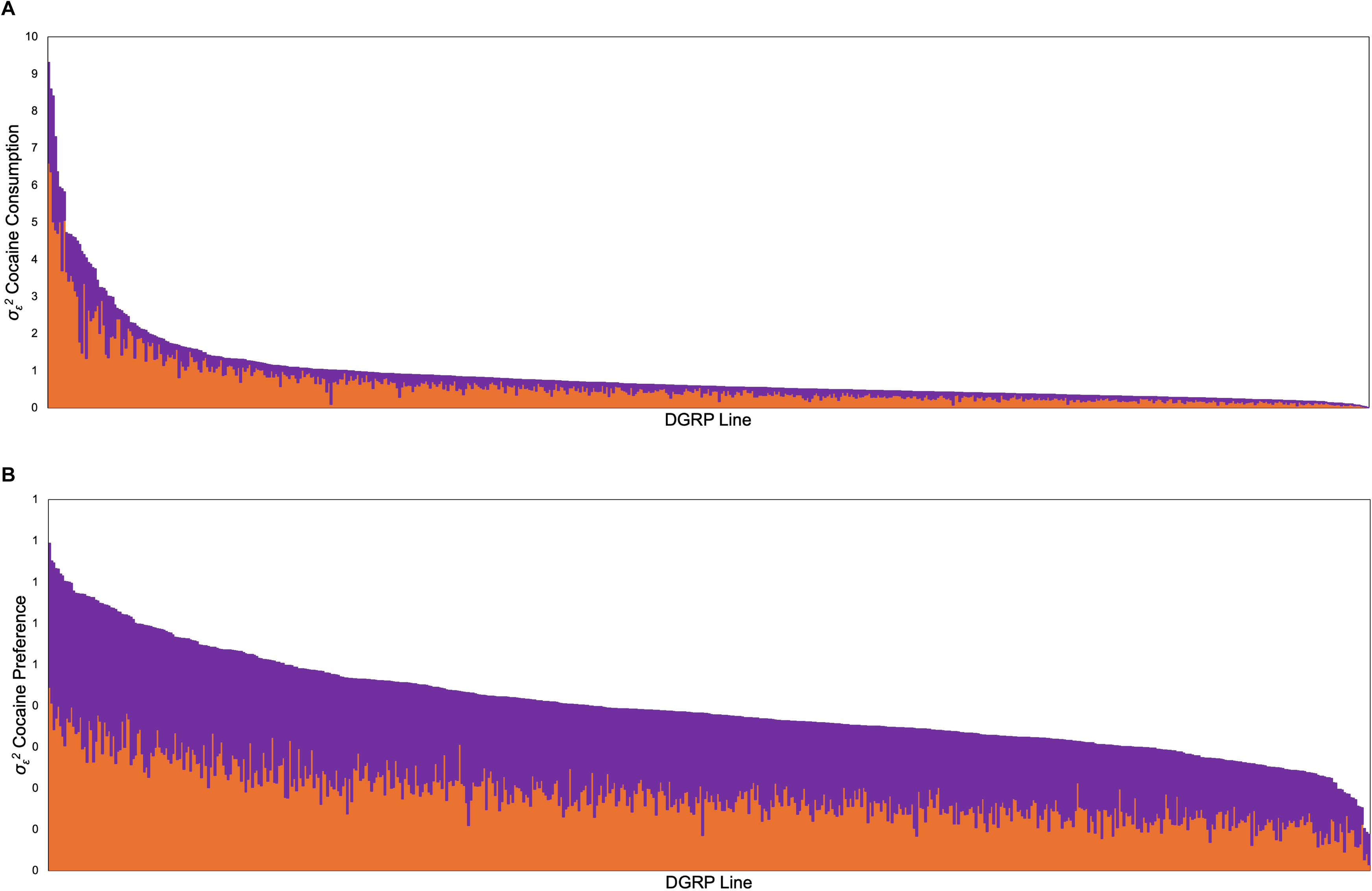
Within line (micro-environmental) variance of cocaine consumption and preference in the DGRP3. (A) Cocaine consumption. (B) Cocaine preference. Stackplots show the estimates of within-line (micro-environmental) variance for each trait in 598 DGRP3 lines. Purple indicates males and orange indicates females.

We computed phenotypic and genetic correlations between the mean and *lnσ_ɛ_* for each trait, for the means of different consumption traits, and for *lnσ_ɛ_* of different traits, separately for males and females (S2D Table). The genetic correlations between the mean and *lnσ_ɛ_* for all traits are very high, implying considerable shared genetic basis of variation for the mean and micro-environmental variation of consumption-related traits. In particular, the genetic correlation between the mean and micro-environmental variation for cocaine consumption is not significantly different from unity in either sex. The genetic correlations between cocaine consumption and preference are high and positive in both sexes, while the genetic correlations between sucrose consumption and cocaine preference are moderately high and negative in both sexes. This is a consequence of the definition of cocaine preference. Although the genetic correlations between mean sucrose and cocaine consumption are low, the genetic correlations between *lnσ_ɛ_* for sucrose and cocaine consumption are high, indicating a shared genetic basis of micro-environmental variation for sucrose and cocaine consumption that is independent of the mean.

### Genome Wide Association (GWA) Analyses

We used the sucrose and cocaine consumption and cocaine preference line means in conjunction with the full genome sequences of the DGRP3 lines to perform GWA analyses to map variants and genes associated with these phenotypes. We performed four GWA analyses for each phenotype (females, males, sex average and sex difference) motivated by our observation that there is significant genetic variation for all traits for each sex separately, for the analyses pooled across sexes, and for the sex by line interaction (modeled by the variance among lines in the sex difference analyses). In addition, we performed GWA analyses using *lnσ_ɛ_* line means for females and males. Note that the broad sense heritabilities of line means for consumption and preference traits approach unity (S2A Table), a favorable scenario for association mapping. These GWA analyses tested for association with ∼2.2 million polymorphic loci with minor allele frequency ≥ 0.01, following adjustment for covariates (*Wolbachia* infection status and 31 large polymorphic inversions, S3 Table) and polygenic relatedness.

We identified 2,155 variants in or near (within 1 kb of the gene body) 866 genes associated with at least one trait at a significance threshold of −*log*_10_(*P*) > 5 (Figures 3, 4, S3-S8 Figs, S4A, S4B Tables). This threshold was chosen based on quantile-quantile plots that show enrichment of −*log*_10_(*P*)-values past this threshold. The only exceptions are for the sex average analysis of sucrose consumption and the *lnσ_ɛ_* analyses for sucrose consumption (S3-S8 Figs). A total of 126 variants in or near 66 genes and 37 intergenic regions have *P*-values less than the conservative 2.3 × 10^−8^ Bonferroni correction for multiple tests (S4C Table). A total of 21 of these genes (*5-HT1A*, *Bsg*, *Ca-beta*, *CanA1*, *CG9664*, *cnc*, *dpr6*, *Exo70*, *fred*, *Hand*, *Irk1*, *Mad*, *nAChRalpha6*, *nkd*, *ptc*, *Ser*, *tkv*, *Toll-7*, *Trim9*, *Trpm*, *vri*) are associated with Gene Ontology (GO) terms involved in nervous system development and function, including behaviors (S4D Table). There are 3,183 site class annotations corresponding to the 2,155 variants associated with one or more consumption traits. The number of annotations exceeds the number of variants because many variants potentially impact more than one gene. Most variants do not affect proteins and are likely regulatory (Table 1).

**Figure 3.**
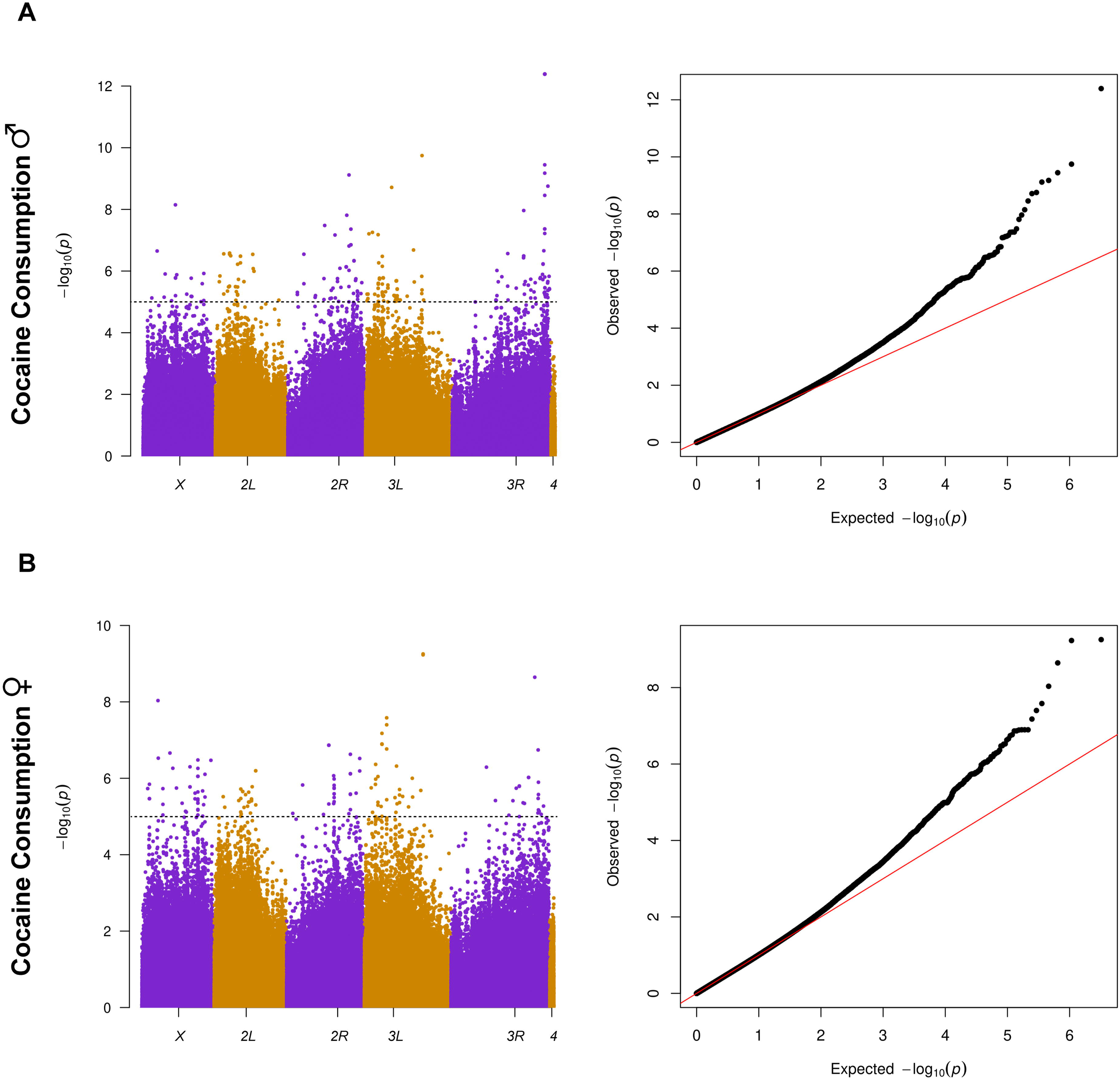
Cocaine consumption GWA results. The left panels show the Manhattan plots, where the *x*-axis denotes the physical location in the *Drosophila* genome with the chromosome arms denoted by color blocks. The *y*-axis is the −*log*_10_(*P*) value for each association test. Each point represents a genetic polymorphism. The horizontal line represents −*log*_10_(*P*) = 5, the nominal significance threshold. The corresponding right panels show the Q-Q plots. (A) Males. (B) Females.

**Figure 4.**
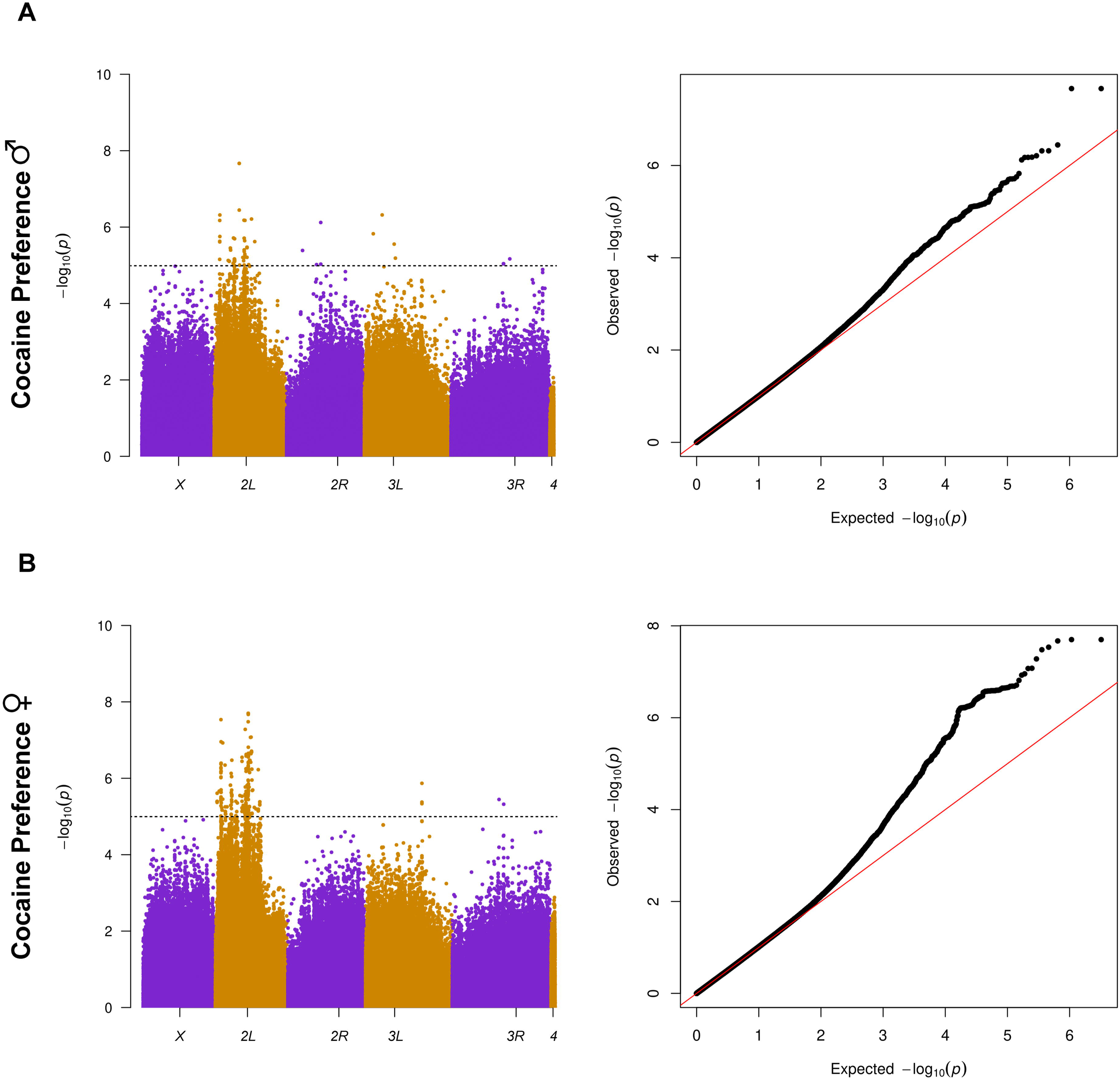
Cocaine preference GWA results. The left panels show the Manhattan plots, where the *x*-axis denotes the physical location in the *Drosophila* genome with the chromosome arms denoted by color blocks. The *y*-axis is the −*log*_10_(*P*) value for each association test. Each point represents a genetic polymorphism. The horizontal line represents −*log*_10_(*P*) = 5, the nominal significance threshold. The corresponding right panels show the Q-Q plots. (A) Males. (B) Females.

**Figure 5.**
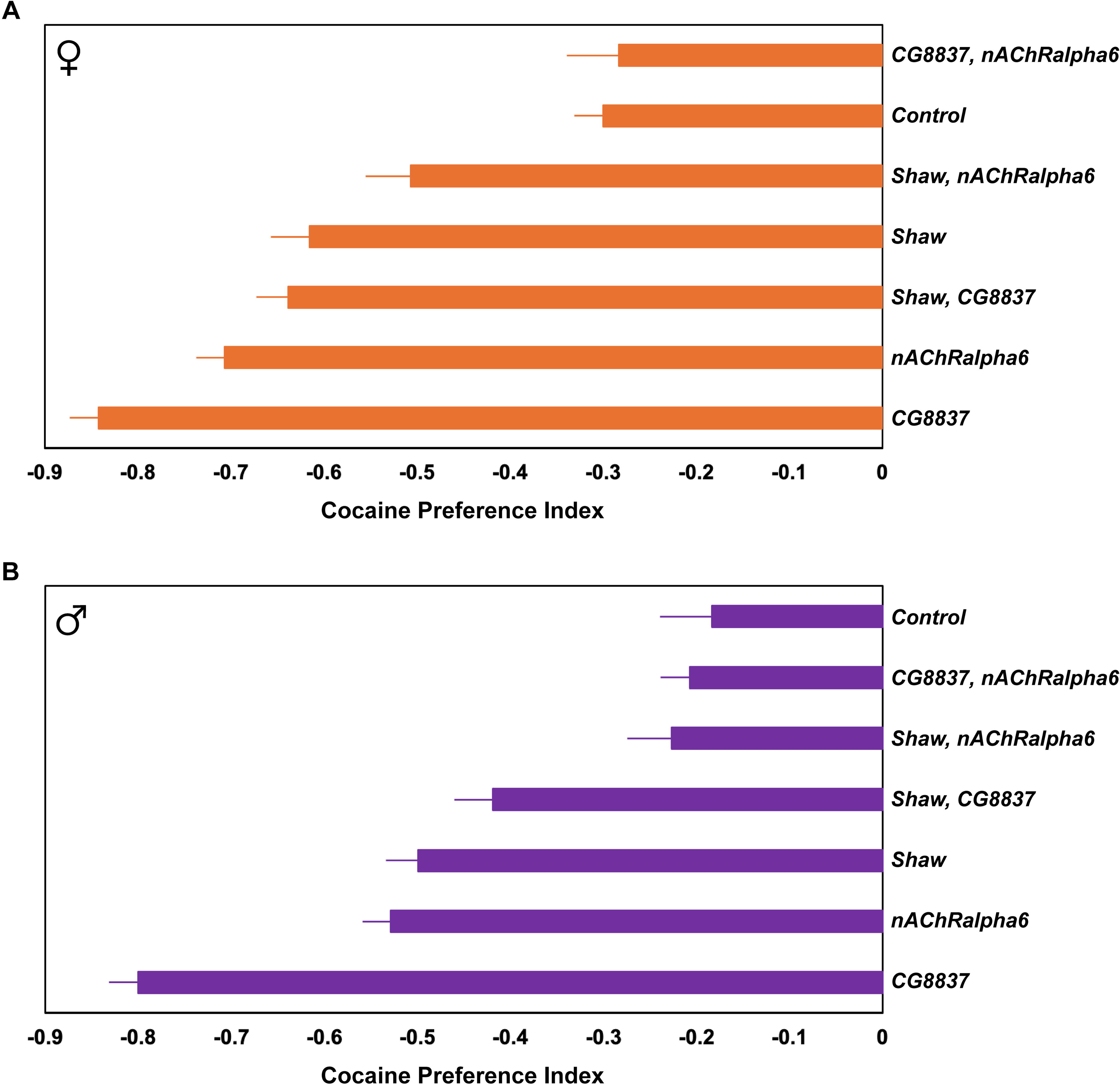
Functional validation of three genetic polymorphisms associated with cocaine preference. Each bar represents the mean cocaine preference index for each indicated genotype. Error bars are standard errors of the mean. The alleles are *2L*_3323727_C for the *CG8837* minor allele, *2L*_3718222_T for the *Shaw* minor allele, and *2L*_9856112_A for the *nAChRalpha6* minor allele. The control genotypes are *2L*_3323727_T, *2L*_3718222_C and 2*L*_9856112_G for *CG8837*, *Shaw* and *nAChRalpha6*, respectively.

**Table 1.** Site Class annotations for variants associated with consumption traits.

| Site Class | Number | Percent |
| --- | --- | --- |
| Intron | 1,090 | 34.244 |
| Intergenic | 524 | 16.463 |
| Upstream | 499 | 15.677 |
| Downstream | 408 | 12.818 |
| Synonymous | 211 | 6.629 |
| 3' UTR | 144 | 4.524 |
| Missense | 109 | 3.424 |
| 5' UTR | 91 | 2.859 |
| Exon, non-coding transcript | 63 | 1.979 |
| Intron, non-coding transcript | 16 | 0.503 |
| Splice polypyrimidine tract, intron | 15 | 0.471 |
| Splice region, splice polypyrimidine tract, intron | 6 | 0.189 |
| Splice donor region, intron | 3 | 0.094 |
| Splice region, synonymous | 2 | 0.063 |
| Splice region, 5' UTR | 1 | 0.031 |
| Splice region, intron | 1 | 0.031 |

We performed GO enrichment analyses [28] using the 779 genes associated with cocaine consumption, cocaine preference, *lnσ_ɛ_* cocaine consumption and/or *lnσ_ɛ_* cocaine preference. Among the top enriched (False Discovery Rate, FDR < 0.05) Biological Process GO terms were nervous system development (GO:0007399) (FDR = 3.89 × 10^−9^), neurogenesis (GO:0022008) (FDR = 4.08 × 10^−9^), neuron differentiation (GO:0030182) (FDR = 4.48 × 10^−9^) and generation of neurons (GO:0048699) (FDR = 4.53 × 10^−9^) (S4E Table). This observation reinforces the inference that variation in genes involved in the development and function of the nervous system that likely affect variation in neural circuitry and connectivity are associated with variation in adult cocaine consumption and preference.

Two other studies [22,23] used the previous version of the DGRP [16,17] to perform GWA analyses of cocaine-related phenotypes. A total of 102 genes identified in this study overlapped with one or both these prior studies, with 86 genes overlapping with Ref. 23 (expected overlap = 35.179, *P* = 9.93 × 10^−15^, hypergeometric test) and 25 overlapping with Ref 22 (expected overlap = 10.37, *P* = 4.29 × 10^−5^, hypergeometric test) (S4F Table). Among these 102 genes, eight (*Ca-beta*, *CG32982*, *Mob2*, *Msp300*, *Prosap*, *pum*, *sif*, *sr*) are present in all three studies; and ten (*Ca-beta*, *cnc*, *fred*, *Msp300*, *ptc*, *Ptip*, *Ser*, *tkv*, *toc*, *Trim9*) are among the 66 genes in this study with variants that had *P*-values less than 2.3 × 10^−8^ (S4F Table). These 102 genes are thus independently validated.

We used the *Drosophila* Integrated Ortholog Prediction Tool (DIOPT) [29] to identify human orthologs of fly genes associated with cocaine consumption, cocaine preference, *lnσ_ɛ_* cocaine consumption and/or *lnσ_ɛ_* cocaine preference. Orthology is defined as a DIOPT composite score ≥ 3. A total of 502 of the 779 genes (64.4%) associated with a cocaine-related phenotype have human orthologs (S4G Table). However, 102 of the 779 genes encode antisense RNAs, long noncoding RNAs, micro RNAs, small nucleolar RNAs and small nuclear RNAs, for which genome sequences are poorly conserved between flies and humans, although analogous human genes may have similar functions. Excluding these genes raises the percentage of human orthologs to 74.2%.

Next, we used DIOPT to determine human functional associations with each ortholog using Online Inheritance in Man, quantitative trait locus mapping and GWA associations (S4G Table). A total of 282 human orthologs of *Drosophila* genes (230 unique human genes and 172 unique fly genes) identified in this study have been associated with addiction phenotypes, substance use disorders, psychiatric disorders and behaviors (S4H Table). We consider all these genes excellent candidates for further study in humans, given the well-known pleiotropy of substance use and psychiatric disorders [30–32]. Six *Drosophila* genes – *Eip63E*, *Rim*, *Fife*, *Dop2R*, *Gad1* and *Rdl* – have human orthologs (*CDK1*, *RIMS2*, *DRD2*, *Gad1*, *Gad2* and *GABRG2*, respectively) associated with cocaine dependence or addiction [3,8,10]. Two human orthologs of *Shaw*, *KCNC1* and *KCNG2*, are associated with opioid sensitivity, while a third (*KCND2*) is associated with alcohol and nicotine co-dependence. Many other human orthologs of fly genes are associated with alcohol dependence (*HPGD*, *SYT17*, *WDR7*, *MICU3*, *ERAP1*), alcohol and nicotine co-dependence (*HTR1A*, *SH3BP5*,), alcohol consumption (*TLR1*), cannabis dependence (*AFF3*), response to amphetamines (*WWOX*, *MAP2K4*), food addiction (*KLHL33*, *NTM*) and eating disorders (*GADL1*, *ABDG1*, *ATP8A2*, *ABOP*, *TRPS1*, *KCNK5*, *MSRA*, *AFF1*, *PDE8A*, *LRP2*, *PPP3CA*, *MCTP1*, *GRID1*, *NTNG1*). A total of 67 unique fly genes with 66 unique human orthologs were associated with sleep phenotypes and 80 unique fly genes with 79 unique human orthologs were associated with schizophrenia and/or bipolar disorder.

### Functional Validation

We used several criteria to prioritize candidate molecular polymorphisms from the GWA analyses for functional validation: a low *P*-value of association with cocaine preference, the DIOPT score [29] for human orthology and the novelty of the association with cocaine preference. We performed out-of-sample validation using DGRP3 lines that were not included in the initial GWA analyses for three molecular polymorphisms: *2L*_3323727 586 bp downstream of *CG8837* (*P* = 4.33 × 10^−8^), *2L*_3718222 in the 5’ UTR of *Shaw* (*P* = 4.84 × 10^−8^), and *2L*_9856112, an intronic polymorphism in *nAChRalpha6* (*P* = 2.39 × 10^−7^). *CG8837* is moderately orthologous to human *SLC2A8*, *SLC2A6* and *SLC2A13* (DIOPT = 3 for each), while *Shaw* and *nAChRalpha6* are highly orthologous (DIOPT = 15) to human *KCNC2/KCNC1* and *CHRNA7/CHRFAM7A*, respectively (S4A, S4G Tables).

We chose 5-6 DGRP3 lines that were not used in the initial GWA analyses that were homozygous for the minor allele of the tested polymorphisms and compared their average cocaine preference to that of 5 control lines that were homozygous for the major alleles for all three polymorphisms. We tested cocaine preference for ∼72 males and ∼72 females for each of these DGRP3 lines, in randomized order, using the same microplate feeder assay as in the original screen (S5A Table). Some of the lines were homozygous for minor alleles at two of the polymorphisms of interest (S5B Table). Therefore, we performed analyses of variance comparing cocaine preference of the control genotypes to that of all minor allele genotypes (S5C Table). The effect of genotype was significant (*P* = 0.0073) for the analyses pooled across sexes as well as for females (*P* = 0.0057) and males (*P* = 0.0143) separately. DGRP3 flies homozygous for the minor allele at only one of the polymorphisms of interest exhibited significantly decreased cocaine preference compared to the control group (Figure 6). However, lines homozygous for minor alleles at two of the polymorphisms of interest had lower cocaine preferences than expected given their single locus effects (Figure 6), suggesting suppressing epistasis.

## Discussion

We assessed the mean and micro-environmental variance of cocaine and sucrose consumption and preference for 598 DGRP3 lines. We found significant genetic variation for all traits, significant genetic variation in sexual dimorphism, and genotype by treatment interaction variance for the consumption traits. Such context-dependent effects are a hallmark of quantitative genetic variation in the DGRP [18,33], including for cocaine-related behaviors [22,23]. Cocaine consumption in *Drosophila* leads to increased incidence of seizures [34]; therefore, preference for cocaine despite adverse physiological effects can be considered an insect analog of human addiction. Our screen identified several DGRP3 lines with innate cocaine preference in at least one sex. Broad sense individual heritabilities of consumption and preference traits were relatively low, ranging from *H*^2^ = 0.102 to *H*^2^ = 0.245. We also observed high positive genetic correlations between consumption and preference traits and micro-environmental variance of consumption and preference traits. This latter observation indicates that high cocaine consumption or cocaine preference genotypes have a large range of cocaine consumption or preference phenotypes, leading to variable penetrance. If this is also true in humans, it could be an additional factor accounting for the difficulty in mapping and replicating variant/gene associations with CUD-related traits.

We performed GWA analyses using line means for all cocaine-related traits. Broad sense heritabilities of line means for consumption and preference that approach unity, the large number of genetically diverse lines used, and the strict control of treatments and other environmental factors together present a favorable scenario for mapping variants and genes associated with cocaine-related traits that is not possible in human populations. Across all traits, we identified 2,155 polymorphisms in/near 866 genes at −*log*_10_(*P*) > 5, a significance threshold for which quantile-quantile plots indicate an excess of true positive associations relative to chance expectation. Further, 126 variants in/near 66 genes and 37 intergenic regions had *P*-values less than the conservative 2.3 × 10^−8^ Bonferroni correction for multiple tests. The genes associated with significant molecular polymorphisms were enriched for GO terms involving all aspects of development, including development and differentiation of the nervous system, and are thus excellent candidate genes affecting cocaine consumption and cocaine preference for future functional studies. The gene encoding the *Gr66a* bitter receptor is associated with the general avoidance behavior of *D. melanogaster* to cocaine [35]. However, genetic variation in *Gr66a* is not associated with the variation we observe in cocaine consumption or preference.

Our GWA analyses replicated 102 genes previously associated with cocaine preference and consumption in *Drosophila* [22,23] using different DGRP lines and an advanced intercross population derived from different DGRP lines. We previously found that Ibrutinib reduces seizures in flies exposed to cocaine [34]. *Btk* is a target for Ibrutinib and an intronic variant in *Btk* was associated with cocaine preference in our GWA analysis. To assess whether the results of the GWA analyses in *Drosophila* have translational potential in humans, we determined known phenotypic associations of human genes orthologous to the fly genes associated with cocaine-related traits. We found six human orthologs of *Drosophila* genes associated with cocaine dependence (*CDK1*, *RIMS2*, *RASGEF1B*, *RBFOX1*, *GPX8, DRD2*) and four human orthologs associated with cocaine addiction (*DRD2*, *GAD1*, *GAD2*, *GABRG2*). Not all human orthologs had associations reaching Bonferroni-corrected significance, suggesting that examining reciprocal ortholog nominally significant associations (−*log*_10_(*P*) > 5) in humans and flies can be an additional tool for identifying plausible candidate genes for further investigation in both species.

Substance abuse traits are often genetically correlated in humans [36–38]; therefore, genes associated with other abused substances may also affect CUD-associated traits but have not been detected to date because sample sizes have been relatively small. In addition to cocaine-related traits, our GWA analyses also identified human orthologs of fly genes associated with opioid sensitivity, alcohol and nicotine co-dependence, alcohol dependence, alcohol consumption, cannabis dependence, response to amphetamines, food addiction and eating disorders. Substance abuse traits are also genetically correlated with other psychiatric disorders: 149 human orthologs of *Drosophila* genes associated with cocaine consumption and preference in our GWA analyses were associated with schizophrenia and/or bipolar disorder, major depressive disorder, autism spectrum disorder, attention deficit hyperactivity disorder, and other psychiatric disorders. In addition, disruptions in sleep and circadian rhythm have been associated with psychiatric disorders, including substance use disorders [39,40], and disrupted sleep or circadian rhythms are risk factors for drug use [41–43]. A total of 66 human orthologs of fly genes associated with cocaine-related traits were associated with sleep phenotypes.

The advantage of having over 1,000 DGRP3 lines is that they can be divided into test and replication sets. We validated three molecular polymorphisms and genes associated with cocaine preference in our initial GWA analysis: *2L*_3323727_T_C near *CG8837*, *2L*_3718222_C_T upstream of *Shaw*, and *2L*_9856112_G_A within *nAChRalpha6*. We observed that DGRP3 lines homozygous for the minor allele of only one of these polymorphisms had low cocaine preference scores relative to the control line, which was homozygous for the major allele at all three loci. However, lines homozygous for the minor allele at two of these loci had increased preference for cocaine, rather than the additive expectation of further reduced preference, implicating suppressing epistasis among these loci.

*CG8837*, which is orthologous to human *SLC2A8*, encodes a transmembrane sugar transporter in *Drosophila* [44], indicating a potential role in modulating glucose uptake and utilization in neuronal cells. Glucose is the primary source of energy for the brain and glucose uptake is essential for synthesis of neurotransmitters [45], synaptic transmission [46], and learning [47]. *Shaw* encodes a voltage-gated potassium channel orthologous to human *KCNC2* [44]. *KCNC2* has been associated with a variety of traits relevant to substance use disorder, including alcohol consumption [48], opioid dependence [49], and lifetime smoking index [50]. Polymorphisms in *KCNC2* impair neuronal excitability, alongside dysregulation of inhibition [51]. *nAChRalpha6* encodes a nicotinic acetylcholine receptor subunit orthologous to human *CHRNA7* [44]. *CHRNA7* has been associated with alcohol consumption [48] and nicotine dependence [52].

In conclusion, our GWA analyses of cocaine-related traits identified novel genes and independently replicated previously identified genes affecting cocaine preference and consumption in *Drosophila*. We have identified *Drosophila* genes with human orthologs associated with cocaine dependence and addiction, which is remarkable considering these genes do not replicate across different human populations and studies. Many of the human orthologs of fly genes implicated by our GWA analyses have been associated with dependence and addiction for other abused substances as well as several psychiatric disorders, consistent with pleiotropic effects observed in humans [30–32,36–38]. We validated the effects of three molecular polymorphisms in three genes associated with cocaine preference in this study with pairwise suppressing epistasis. Positive correlations between mean cocaine consumption and micro-environmental variance of cocaine consumption combined with suppressing epistasis reduce the effects of molecular polymorphisms associated with cocaine-related traits. These results demonstrate the conservation of genetic factors influencing cocaine-related traits across species and highlight the translational power of forward genetic studies in *Drosophila*.

## Methods

### D. melanogaster Lines

We used 598 DGRP3 lines and Canton S (B) (CSB) control flies. We reared all flies on molasses/cornmeal/yeast culture medium at 25°C, a 12-hour light-dark cycle, and 50% humidity. We placed the parents of the DGRP3 flies used for consumption assays in controlled adult density bottles containing 20 males and 20 females, and the parents of CSB flies used for consumption assays in controlled adult density vials containing 5 males and 5 females. We quantified cocaine and sucrose consumption and cocaine preference for adult progeny aged 3-5 days post-eclosion.

### Quantification of Consumption and Preference

We anesthetized the flies used to quantify consumption and preference using CO_2_ and sorted them into individual wells of a 96-well microplate containing 80 μL of 1.5% agar medium per well (to prevent desiccation). We used two microplate replicates for each DGRP3 line, with each replicate containing 36 DGRP3 males, 36 DGRP3 females, 6 CSB control males, and 6 CSB control females. We included evaporation control wells with each replicate to account for loss of liquid during the exposure period, and CSB controls to account for variation among replicate microplates. We covered the microplates with a 3D-printed coupler device [25]. We deprived the flies of food for 6 hours to encourage consumption and allow excretion of solid material that could interfere with spectrophotometric measurements of consumption. Following this food deprivation period, we allowed the flies to feed *ad libitum* with a choice between control and 0.02% cocaine-supplemented solutions.

We made the two solutions used to quantify preference and consumption fresh daily. The control solution contained 4% sucrose, 1% yeast extract, and 40 μg/mL of FD&C Blue #1; and the cocaine solution contained 4% sucrose, 1% yeast extract, 0.2 mg/mL cocaine and 40 μg/mL of FD&C Blue #1. We obtained cocaine-HCl from NIDA under DEA license RA0443159. We exposed the flies to 10 μL of each solution for a period of 22 hours in the microtiter plates and measured the absorbance values of each solution using a SpectraMax iD5 plate reader and SoftMax Pro 7.1 software before and after consumption [25]. We indexed plate replicates before post-exposure measurements to account for deceased flies, which were omitted from further analyses. We translated the absorbance values to volumes using a custom R script (https://github.com/jwalte5/mfa96/). We adjusted raw absorbance values for within-replicate evaporation and between replicate variation using evaporation control wells and CSB control flies, respectively. We obtained cocaine preference values for each fly from the volumes of sucrose and cocaine consumed as: Preference Index = (Volume of cocaine - Volume of sucrose) / (Volume of cocaine + Volume of sucrose). This index normalizes for the total volume that each fly consumes, delineating cocaine preference from variation in overall consumption patterns across the DGRP3 lines.

### Quantitative Genetic Analyses

We performed data processing in R [53], and statistical modeling using SAS [54]. We assessed differences in consumption using a three-way factorial mixed effects analysis of variance (ANOVA): *Y* = *μ* + *S* + *L* + *T* + *S* × *L* + *S* × *T* + *L* × *T* + *S* × *L* × *T* + *Rep*(*T* × *L*) + *S* × *Rep*(*T* × *L*) + *ε*, where *Y* is volume consumed, *μ* is the overall mean, *S* is sex (fixed), *T* is treatment (fixed effect of solution (control and cocaine)), *L* is DGRP3 line (random), *Rep* is microplate replicate (random), and *ε* is the residual error. We also performed reduced analyses by sex, by treatment, and by sex and treatment. The full ANOVA model for preference is a two-way factorial ANOVA: *Y* = *μ* + *S* + *L* + *S* × *L* + *Rep*(*L*) + *S* × *Rep*(*L*) + *ε*, where the terms are defined above. We estimated broad sense heritability (*H*^2^) from the full consumption model as 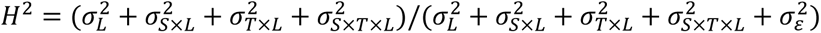, where *σ*^2^ denotes a variance component for the random effects indicated by the subscripts [27]. Similarly, we estimated broad sense heritabilities for female consumption 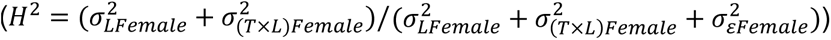, male consumption 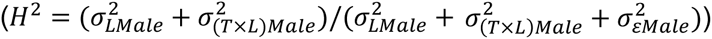, sucrose consumption 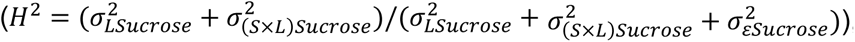, cocaine consumption 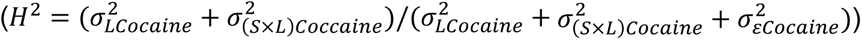, cocaine preference 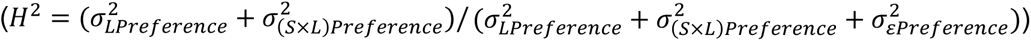 and all traits reduced by sex and treatment 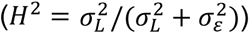. We computed genetic correlations of consumption traits between males and females (*r_GS_*) as 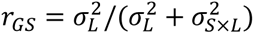, between treatments (*r_GT_*) as 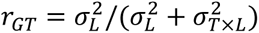, and, for the full consumption model, across sexes and treatments (*r_GST_*) as 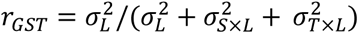.

We evaluated whether there was heterogeneity in micro-environmental variance (within-line variance, 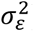) using Levene’s Test [55]. We estimated the broad sense heritability of micro-environmental variance by computing *ln*(*σ_ε_*) for each microplate replicate for each trait [26]. Heritability estimates from the full models are 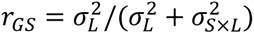 and from the reduced models are 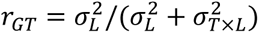. We also computed the cross-sex genetic correlations of micro-environmental variance as 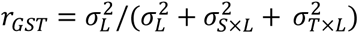.

We estimated cross-trait phenotypic correlations using Pearson product moment correlation coefficients of line means. We estimated pairwise cross-trait genetic correlations between traits X and *Y* (*r_GXY_*) as *r_GXY_* = *Cov_XY_*/*σ_LX_σ_LY_*, where *Cov_XY_* is the covariance between the traits (numerator of Pearson’s correlation coefficient) and *σ_LX_* and *σ_LY_* are the square roots of the line variance components for traits X and *Y* [27]. We estimated cross-trait phenotypic and genetic correlations separately for males and females. We computed standard errors of the correlations 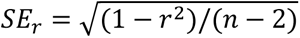 [56].

### Genome-Wide Association Analyses

We performed single variant tests of association for ∼2.2 million polymorphic variants with minor allele frequencies ≥ 0.01. We performed these GWA analyses for mean cocaine consumption, sucrose consumption and cocaine preference using line means for females, males, the average of females and males and the difference between the sexes. We also performed GWA analyses for micro-environmental variance (*lnσ_ε_*) of cocaine consumption, sucrose consumption and cocaine preference for males and females. We filtered PLINK files and covariate files (*Wolbachia* infection status and 31 large polymorphic inversions) for the DGRP3 lines that were used and sites with a genotype missing rate ≤ 0.1. We determined the covariates to be included into the linear mixed effects modeling (LMM) for association testing by fitting a generalized linear model for each covariate against the phenotype. We included covariates with *P*-values less than 0.1. We also included the Genomic Relationship Matrix (GRM), calculated using GEMMA v0.98.5 [57], as a covariate to account for cryptic relatedness. We performed the GWA analyses using GEMMA and used the Wald test to determine significance. The β coefficients from the Wald test are estimates of additive effects, calculated as half the difference between means of individuals with the major and minor allele genotypes [27]. We generated quantile-quantile (Q-Q) plots and Manhattan plots in R using qqman library [58]. The Q-Q plots showed enrichment for *P*-values ≤ 10^−5^, which we used as the significance threshold. We annotated genes associated with genetic polymorphisms within 1 kb of the gene body using Variant Effect Predictor v111 [59] and genes from FlyBase [44]. We performed Gene Ontology (GO) enrichment and pathway analyses using PANTHER [28,60]. We used the *Drosophila* Integrated Ortholog Prediction Tool (DIOPT) [29] to identify human orthologs of fly genes associated with cocaine-related phenotypes with a DIOPT score ≥ 3, and to determine human functional associations with each ortholog.

### Functional Validation

We selected DGRP3 lines that were not previously tested for cocaine preference for out-of-sample validation of three polymorphisms identified in the genome-wide association analysis. DGRP3 lines homozygous for the minor allele of interest were selected at position *2L*_3323727 for validation of *CG8837* (*n* = 5), *2L*_3718222 for *Shaw* (*n* = 6), and *2L*_9856112 for *nAChRalpha6* (*n* = 5). Some of the lines were homozygous for minor alleles at two of the polymorphic sites. We also selected Control lines (*n* = 5) containing the major allele at all three of these positions from the previously untested DGRP3 lines. We reared these lines and quantified cocaine preference exactly as described above for the initial screen. We analyzed the preference data using the following mixed effects factorial ANOVA model: *Y* = *μ* + *G* + *S* + *G* × *S* + *L*(*G*) + *S* × *L*(*G*) + *Rep*(*G* × *L*) + *S* × *Rep*(*G* × *L*) + *ε*, where *G* indicates the fixed effect of genotype (control lines with major alleles at all three polymorphic sites versus lines containing the minor allele at one or more sites) and all other terms are as defined above.

## Supplementary Information

**S1 Figure. Variation in sucrose consumption in 598 DGRP3 lines.** The circles represent line means for males (purple, N = 36,731) and females (orange, N = 38,144) and error bars are standard errors of the mean. The line means are ordered by increasing male sucrose consumption.

**S2 Figure. Micro-environmental variance of sucrose consumption in 598 DGRP3 lines.** Stackplots show the estimates of within-line (micro-environmental) variance. Purple indicates males and orange indicates females.

**S3 Figure. Cocaine consumption GWA results for the average of males and females and the difference between the sexes.** The left panels show the Manhattan plots, where the *x*-axis denotes the physical location in the *Drosophila* genome with the chromosome arms denoted by color blocks. The *y-*axis is the −*log*_10_(*P*) value for each association test. Each point represents a genetic polymorphism. The horizontal line represents −*log*_10_(*P*) = 5, the nominal significance threshold. The corresponding right panels show the Q-Q plots. (A) Sex average. (B) Sex difference.

**S4 Figure. Cocaine preference GWA results for the average of males and females and the difference between the sexes.** The left panels show the Manhattan plots, where the *x*-axis denotes the physical location in the Drosophila genome with the chromosome arms denoted by color blocks. The *y*-axis is the −*log*_10_(*P*) value for each association test. Each point represents a genetic polymorphism. The horizontal line represents −*log*_10_(*P*) = 5, the nominal significance threshold. The corresponding right panels show the Q-Q plots. (A) Sex average. (B) Sex difference.

**S5 Figure. Sucrose consumption GWA results.** (A) Males. (B) Females. (C) Sex average. (D) Sex difference. The left panels show the Manhattan plots, where the *x*-axis denotes the physical location in the *Drosophila* genome with the chromosome arms denoted by color blocks. The *y*-axis is the −*log*_10_(*P*) value for each association test. Each point represents a genetic polymorphism. The horizontal line represents −*log*_10_(*P*) = 5, the nominal significance threshold. The right panels show the corresponding Q-Q plots.

**S6 Figure. *lnσ_ε_* cocaine consumption GWA results.** The left panels show the Manhattan plots, where the *x*-axis denotes the physical location in the *Drosophila* genome with the chromosome arms denoted by color blocks. The *y*-axis is the −*log*_10_(*P*) value for each association test. Each point represents a genetic polymorphism. The horizontal line represents −*log*_10_(*P*) = 5, the nominal significance threshold. The corresponding right panels show the Q-Q plots. (A) Males. (B) Females.

**S7 Figure. *lnσ_ε_* cocaine preference GWA results.** The left panels show the Manhattan plots, where the *x*-axis denotes the physical location in the *Drosophila* genome with the chromosome arms denoted by color blocks. The *y*-axis is the −*log*_10_(*P*) value for each association test. Each point represents a genetic polymorphism. The horizontal line represents −*log*_10_(*P*) = 5, the nominal significance threshold. The corresponding right panels show the Q-Q plots. (A) Males. (B) Females.

**S8 Figure. *lnσ_ε_* sucrose consumption GWA results.** The left panels show the Manhattan plots, where the *x*-axis denotes the physical location in the *Drosophila* genome with the chromosome arms denoted by color blocks. The *y*-axis is the −*log*_10_(*P*) value for each association test. Each point represents a genetic polymorphism. The horizontal line represents −*log*_10_(*P*) = 5, the nominal significance threshold. The corresponding right panels show the Q-Q plots. (A) Males. (B) Females.

**S1 Table. Raw data and line means.** (A) Raw cocaine and sucrose consumption data. (B) Mean and micro-environmental variance 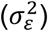 of consumption and preference traits in the DGRP3.

**S2 Table. Quantitative genetic analyses.** (A) Three-way factorial mixed effects ANOVA for consumption and reduced analyses, and two-way factorial mixed effects ANOVA for preference and reduced analyses. ANOVA terms are sex (*S*, fixed), treatment (*T*, sucrose or cocaine solution, fixed), DGRP3 line (*L*, random), all interactions of main effects, plate replicate (*Rep*, random) and residual (*E*, error). df: degrees of freedom. MS: Type III mean squares. *σ*^2^: variance component. (B) Levene tests for heterogeneity of environmental variance. (C) Two-way factorial mixed effects ANOVA for micro-environmental variance of consumption, preference and reduced analyses. (D) Phenotypic and genetic correlations.

**S3 Table. *P*-values for association of covariates with consumption and cocaine preference used in the GWA analyses.** Covariates with *P*-values < 0.1 (highlighted cells) were included in the GWA analysis for each trait. Inversions are denoted by the chromosome arm, start breakpoint position (P) and the inversion length (L), both in base pairs.

**S4 Table. Compiled GWA analyses for consumption and preference traits.** (A) GWA analyses of variants and genes significantly (*P* < 10^−5^) associated with cocaine consumption, sucrose consumption, cocaine preference, and micro-environmental variance of these traits. The GWA analyses for the consumption and preference traits were performed for females, males, the average of the two sexes and the difference between sexes. The GWA analyses for the micro-environmental variances were only performed for males and females. *P*-values in red font are less than the Bonferroni significance threshold. Colored cells for Trait and Analysis denote the same variants associated with different traits or analyses. (B) All associated variants and genes. (C) GWA results for Bonferroni-significant variants. (D) Gene Ontologies for Bonferroni-significant genes. (E) Gene Ontology enrichment for associated genes. (F) Overlap of associated genes with previous studies. (G) Human orthologs of associated fly genes and their associations with human diseases. (H) Associated fly genes and their human orthologs associated with psychiatric disorders and behavioral traits.

**S5 Table. Functional validation analyses.** (A) Raw data. (B) Line means. (C) Two-way factorial mixed effects ANOVAs for cocaine preference and reduced analyses by sex. ANOVA terms are sex (*S*, fixed), genotype (*G*, fixed), DGRP3 line (*L*, random), interactions of main effects, plate replicate (*Rep*, random) and residual (*E*, error). df: degrees of freedom. MS: Type III mean squares.

## Acknowledgments

This work was supported by National Institutes of Health grants F31 DA057062-02 to J.S.H. and U01 DA041613 and P20 GM139769 to T.F.C.M. and R.R.H.A.

