## Supplementary figures and images for "Genetics of Cocaine Consumption and Preference in *Drosophila melanogaster*"

### Figure S1

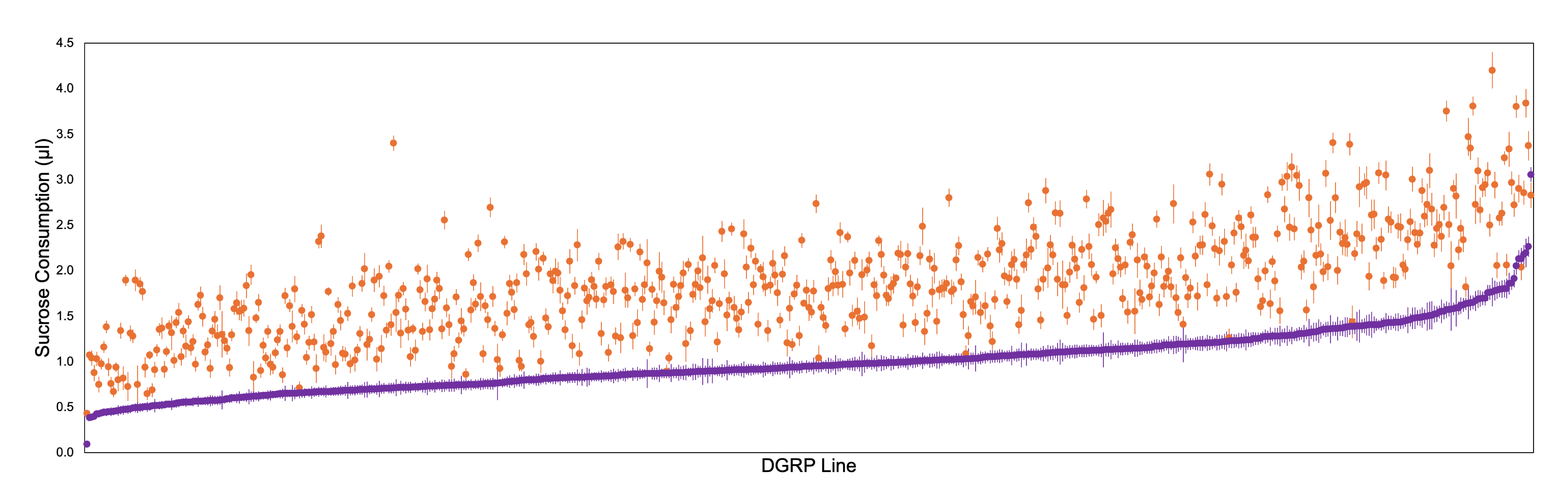

### Figure S2

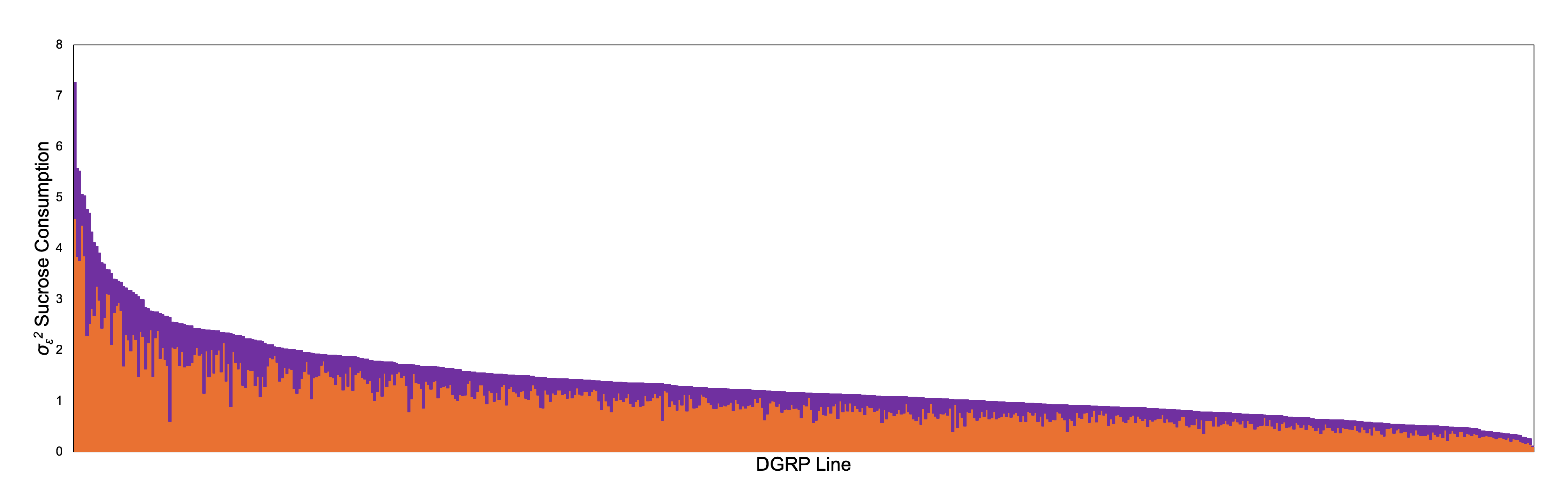

### Figure S3

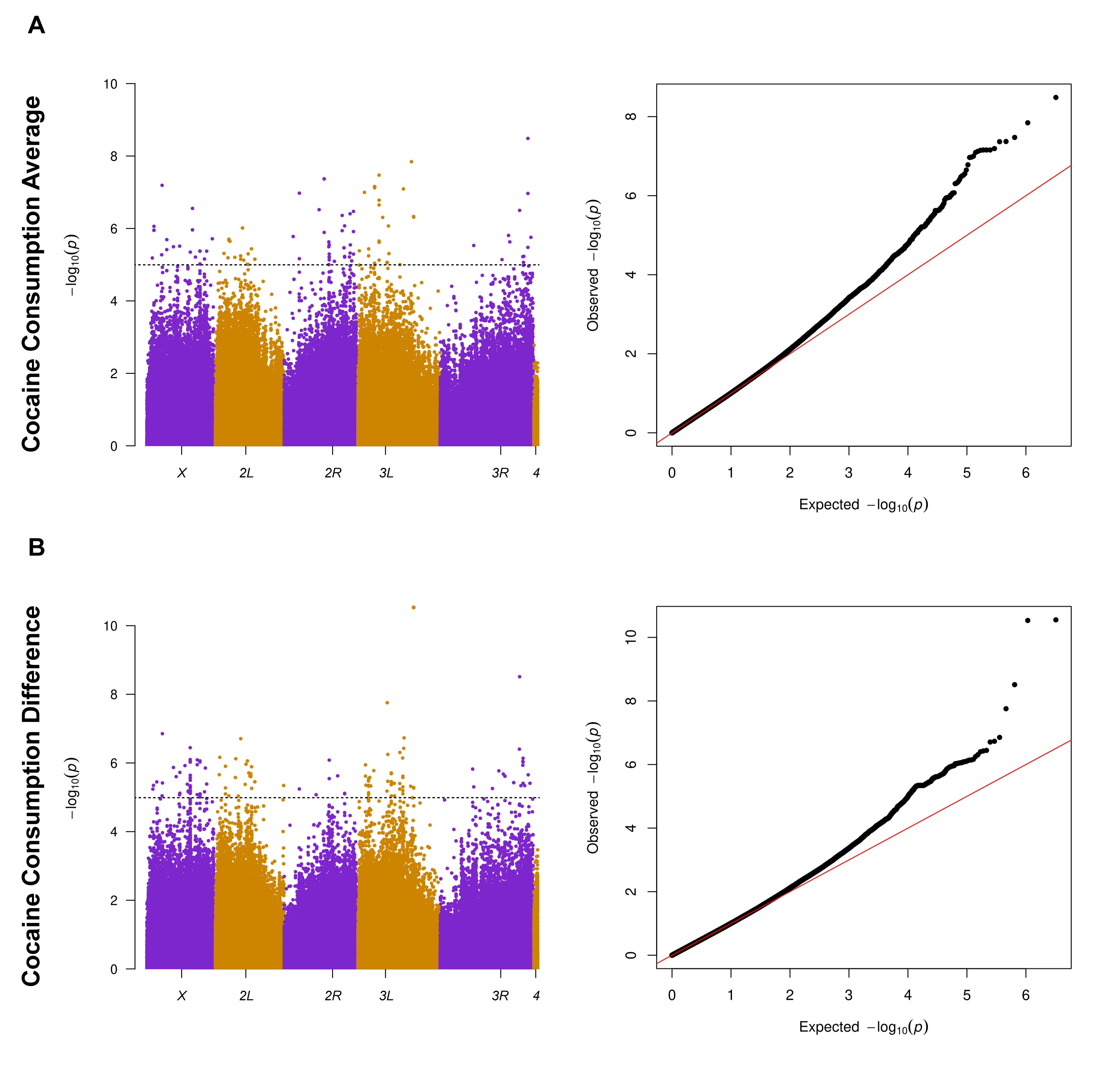

### Figure S4

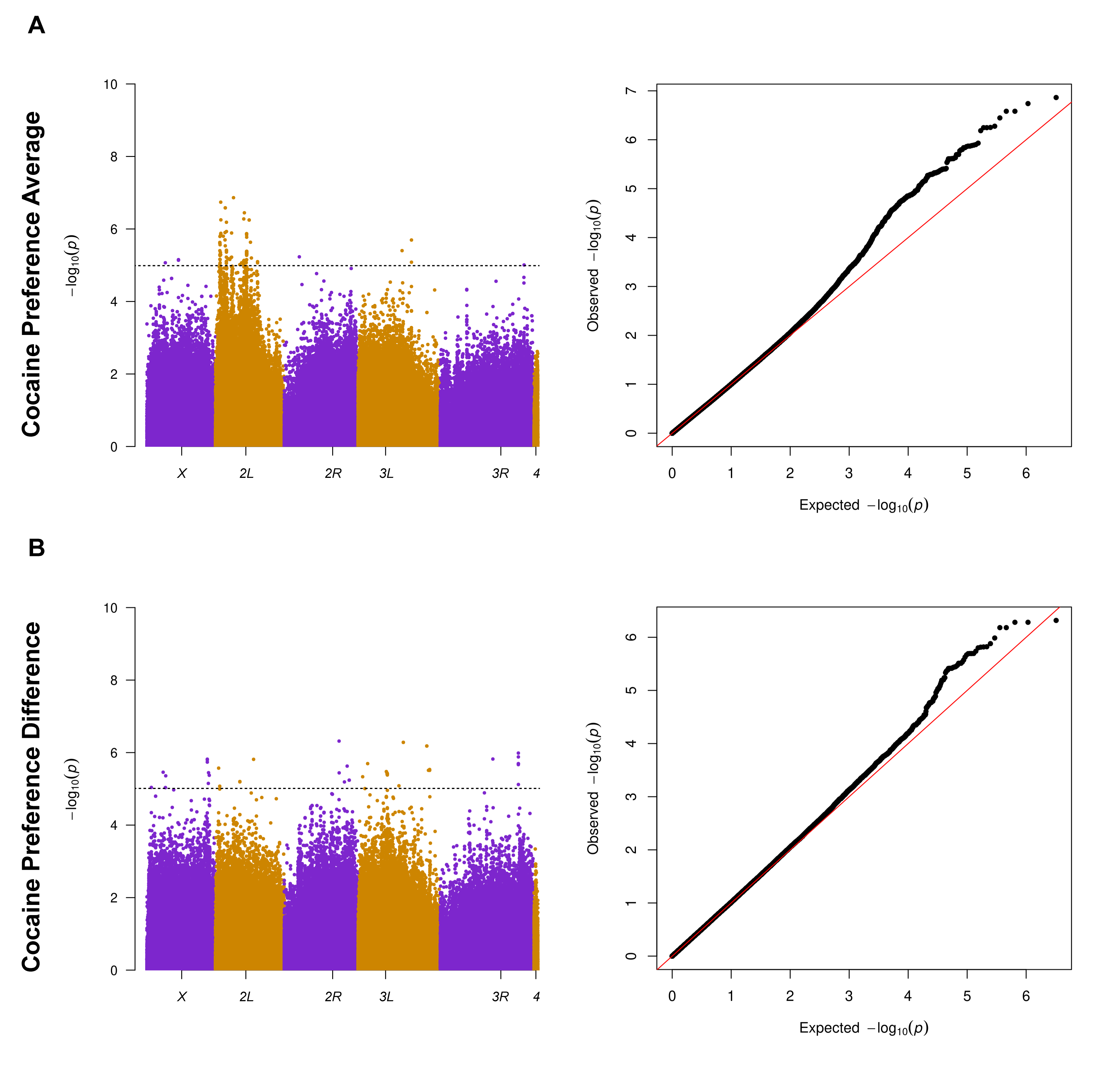

### Figure S5

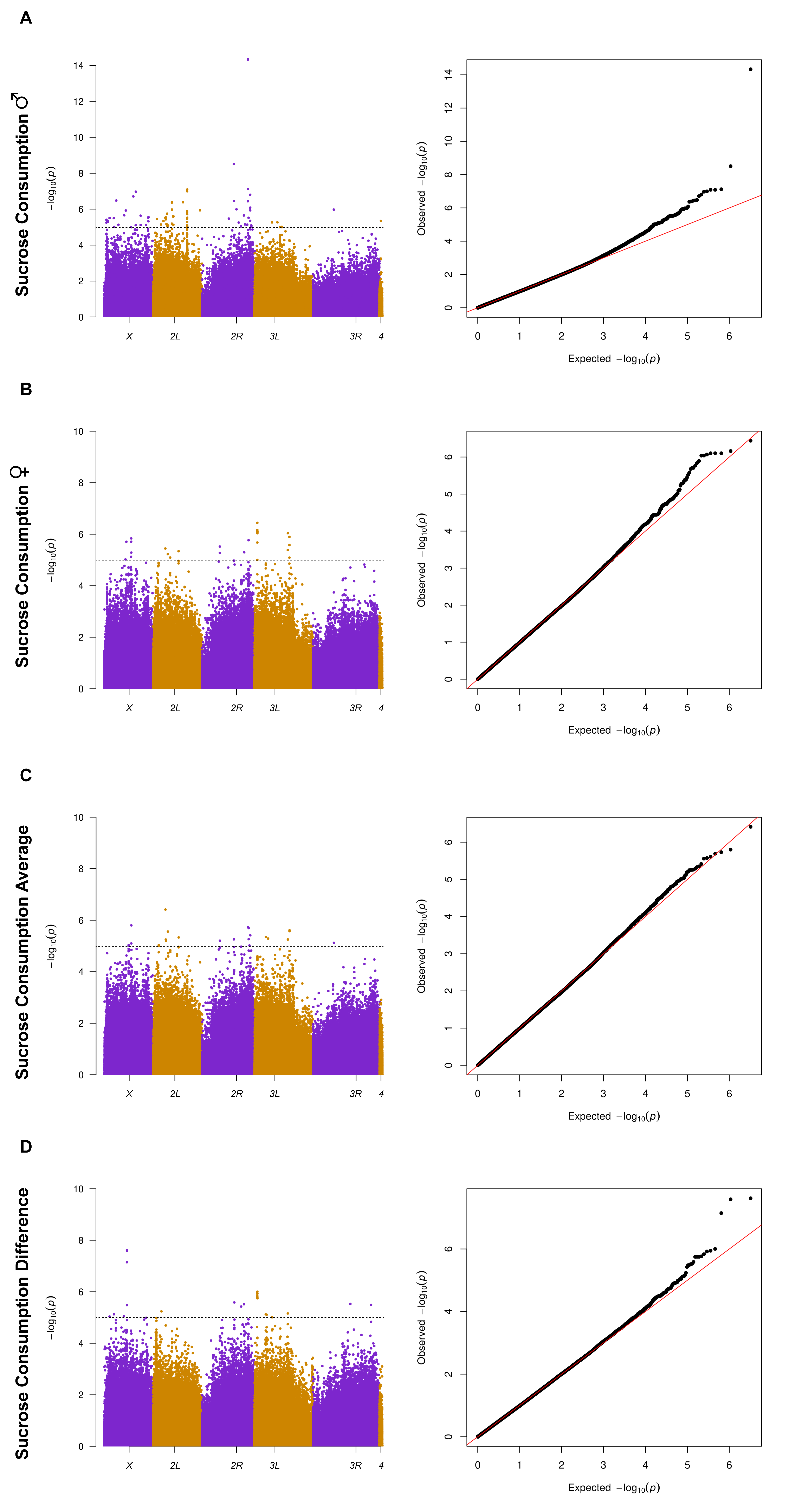

### Figure S6

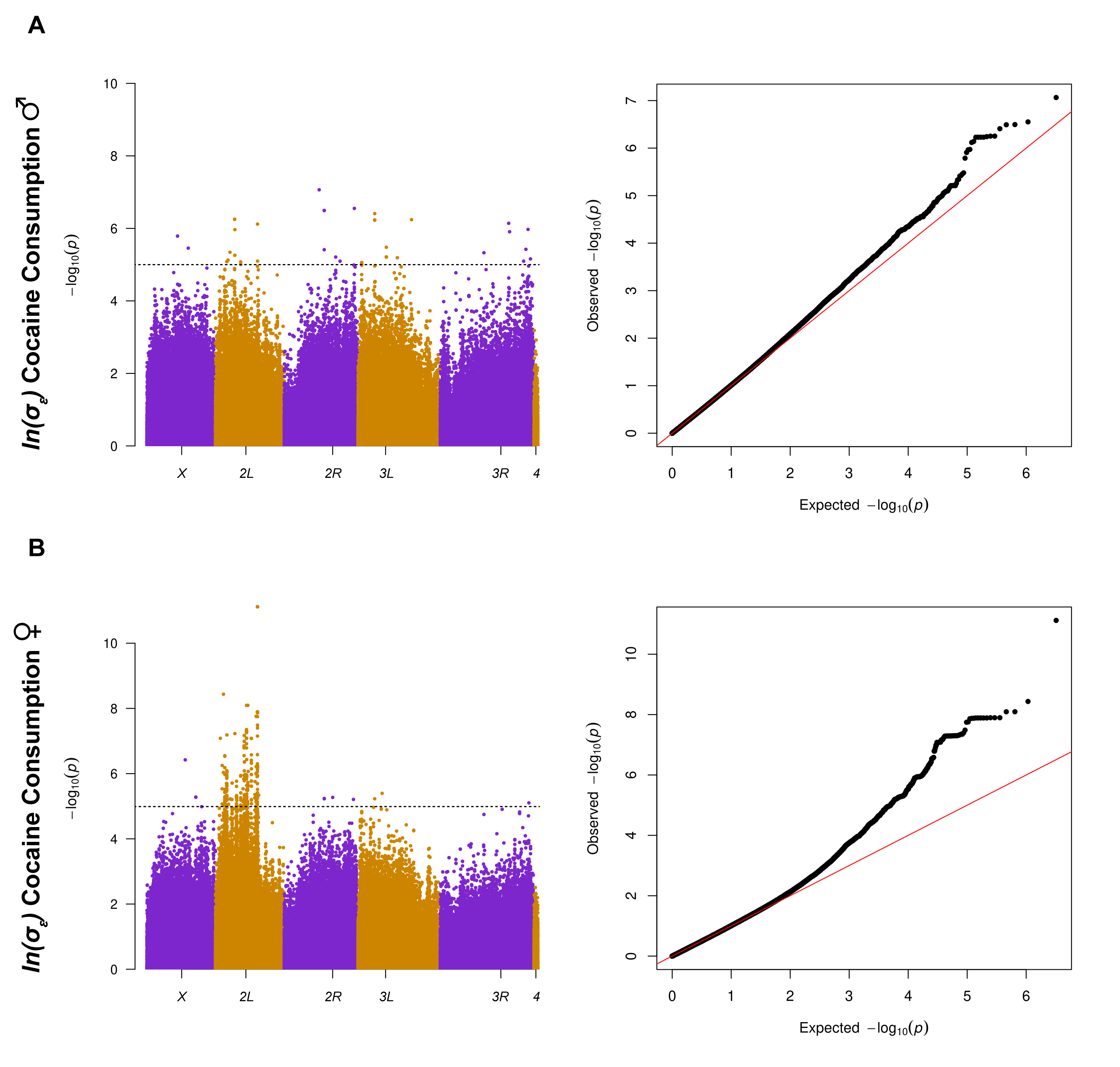

### Figure S7

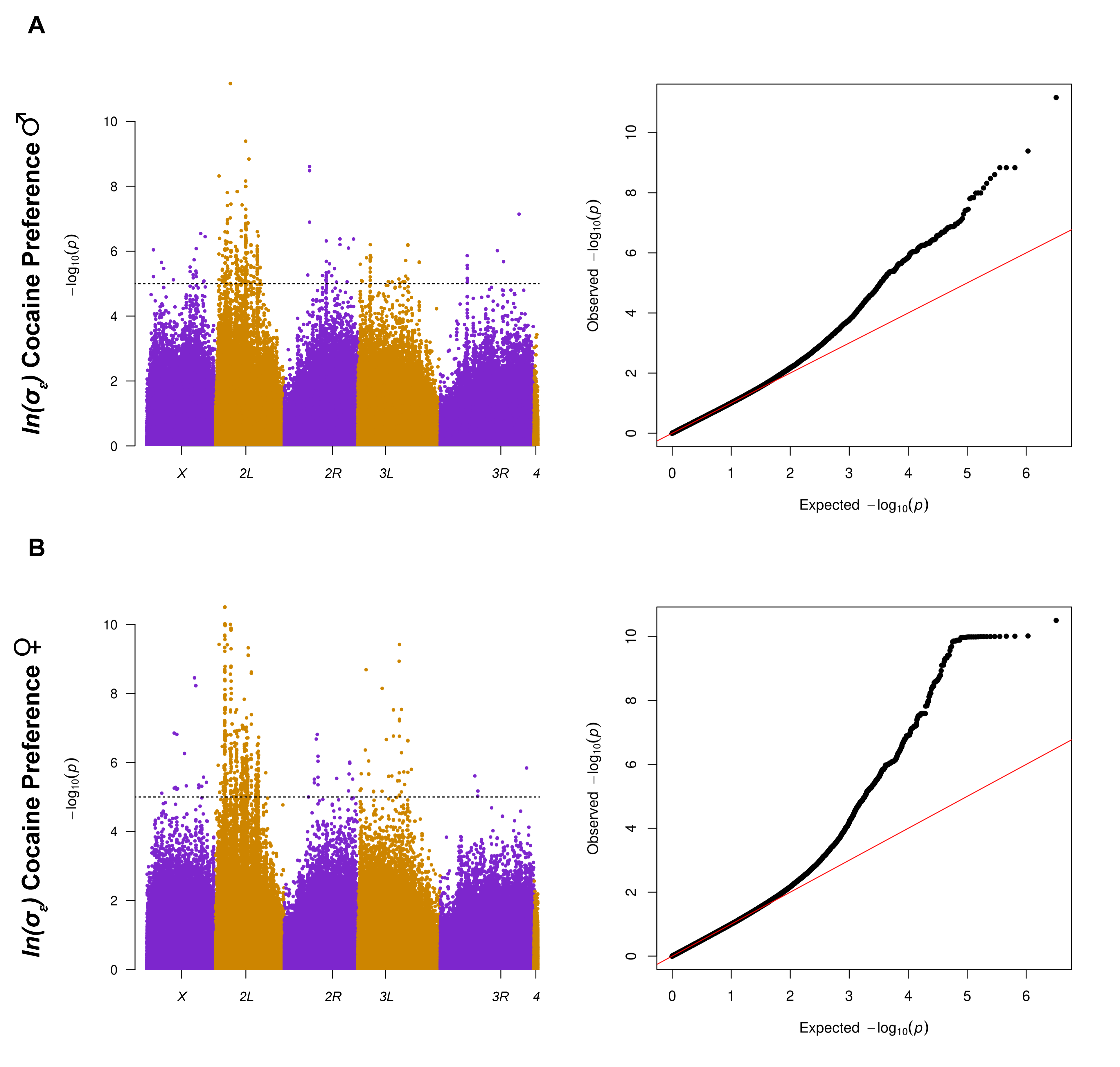

### Figure S8

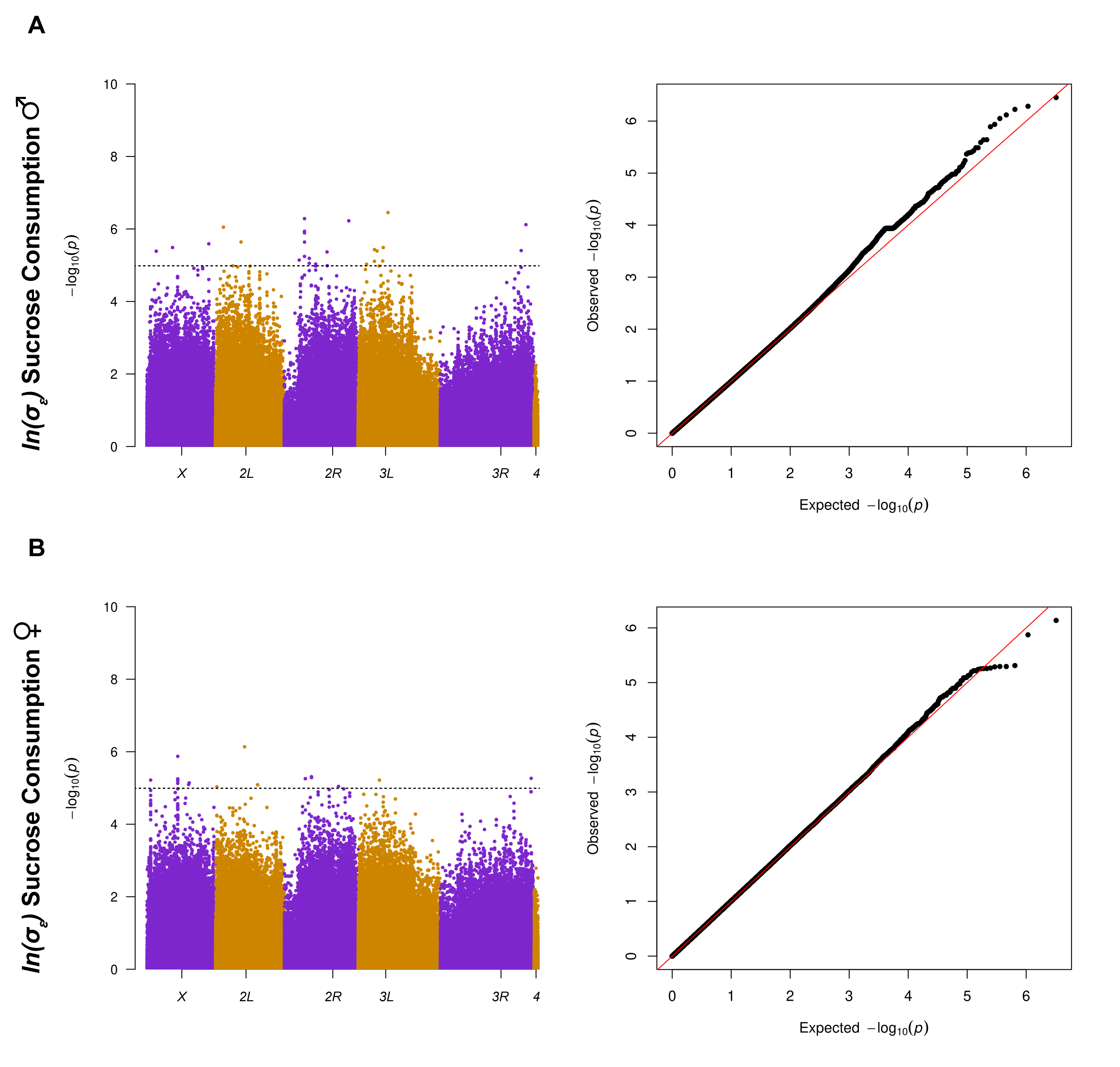
